# Lung progenitors’ exhaustion in response to microenvironment stress during aging

**DOI:** 10.64898/2026.02.18.706624

**Authors:** Fernanda Toscano-Marquez, Ángeles Garcia-Vicente, Uriel Camacho-Silverio, Tania Valdivia-Herrera, Mariana Río de la Loza, Elisa Hernández-Xóchihua, Remedios Ramírez, Moises Selman, Annie Pardo, Yair Romero

**Author notes:** Correspondence should be addressed to: Yair Romero, PhD, Facultad de Ciencias, Universidad Nacional Autónoma de México, Av. Universidad 3000, Coyoacán 04510, CDMX, México.

## Abstract

Progenitor cells in aged tissues undergo changes in their microenvironment that may impact their functionality during regeneration. Despite recent advances in understanding the role of adult lung progenitors, the impact of aging on these cells remains unclear. To analyze aging modifications, we used aged wild-type mice of 18-24 months old, and Zmpste24^-/-^ deficient mice, which exhibit an accelerated aging phenotype. A three-dimensional organoid culture system was employed to assess the lung regeneration capacity. Additionally, mouse epithelial cells and fibroblasts were isolated and characterized with senescence and autophagy markers. Our findings revealed that lung epithelial cells from aged mice and Zmpste24^-/-^ mice hold their regeneration capacity, maintaining their phenotype and a healthy cellular state through an increase in autophagy, particularly when co-cultured with healthy fibroblasts. Conversely, cultured fibroblasts from Zmpste24^-/-^ mice show nuclear defects and acquire a senescent phenotype, characterized by mTORC1 activation and reduced autophagy, which in turn impairs organoid formation. Moreover, these progenitor cells become increasingly susceptible to mechanical stress with aging due to reduced nuclear lamins and the Zmpste24 defect. This vulnerability is illustrated by FACS sorting, which can further compromise their regenerative potential. Our results indicate that, in aging, progenitor cells and their fibroblast niche integrate microenvironmental signals that shape cell-cell interactions essential for lung regeneration.

## Introduction

The aging phenotype is associated with a limited regenerative capacity, mainly due to stem cell exhaustion. The lung is considered a low-turnover organ, where regeneration depends on the activity of rare stem cells and facultative progenitors [1]. In the airway epithelium, basal cells have been identified as rare stem cells responsible for its regeneration [2]. In the alveolar region, regeneration is primarily mediated by facultative progenitors, particularly a subpopulation of alveolar epithelial type 2 (AT2) that can restore alveolar structure through the interaction with a single-cell niche composed of fibroblasts [3–5]. Crucially, in the alveolar epithelium such cell–cell interactions influence cell fate decisions [6]. Considering the central importance of such interactions in orchestrating regenerative processes, it is therefore essential to determine whether alterations associated with aging prevent lung regeneration and promote stem cell exhaustion.

Despite recent advances in understanding adult lung stem cells, the effect of aging on their function remains largely unknown. To address this gap, we compare physiological and accelerated aging using three distinct mouse models: Zmpste24-deficient mice (a model of premature aging), physiologically aged wild-type mice, and young wild-type mice controls. Zmpste24 is a metalloprotease critical for lamin A/C processing. Although Zmpste24-deficient mice do not exhibit an aging phenotype at early stages, by 16 weeks they display hallmark features of premature aging, such as weight loss, graying hair, and reduced lifespan compared to wild-type counterparts [7,8]. This model thus provides a valuable tool to explore whether accelerated aging recapitulates key aspects of physiological lung aging.

Lopez-Otín, *et al*. have proposed that among the hallmarks of aging, stem cell exhaustion and altered intercellular communication are key determinants of the aging phenotype. Although all hallmarks are interconnected, they can be conceptually organized as a sequence of events. Initially, primary features emerge, such as DNA damage, epigenetic changes, loss of proteostasis and impaired autophagy [9,10]. Among these, defective autophagy is particularly critical, as it comprises stem cells function and survival [11]. Autophagy maintains cellular homeostasis by removing damage or unnecessary cytoplasmic components through the lysosomal pathway [12].

In later stages, certain responses act as defense mechanisms but may become detrimental when dysregulated. These include antagonistic processes such as cellular senescence and autophagy since in the context of aging, autophagy prevents cellular senescence by preserving stem cell function and viability promoting regeneration [13–15]. The mTOR pathway serves as a central regulator, integrating signals to modulate stem cell potential in a tissue-specific manner [16].

Therefore, the aim of this study was to analyze the potential role of autophagy and senescence within the regenerative niche, to determine whether age-related alterations in these processes contribute to AT2 progenitor cell exhaustion and impaired lung regeneration.

## Results

### Progenitor epithelial cells of aged mice hold the regenerative potential

To evaluate the capacity of lung regeneration, epithelial cells were cultured in organoids, a model that allows its evaluation *in vitro* [17]. In brief, epithelial cells isolated from mice were mechanically and enzymatically dissociated and positively selected with the EpCam marker by magnetic beads (MACS) (**Fig. 1a**). This procedure yielded a population enriched in epithelial cells, in which the majority displayed a type II phenotype, as validated by the expression of surfactant protein C (Sftpc) through RT-qPCR and proSPC immunostaining **(Fig. 1b and c)** [4,5].

**Fig. 1.**
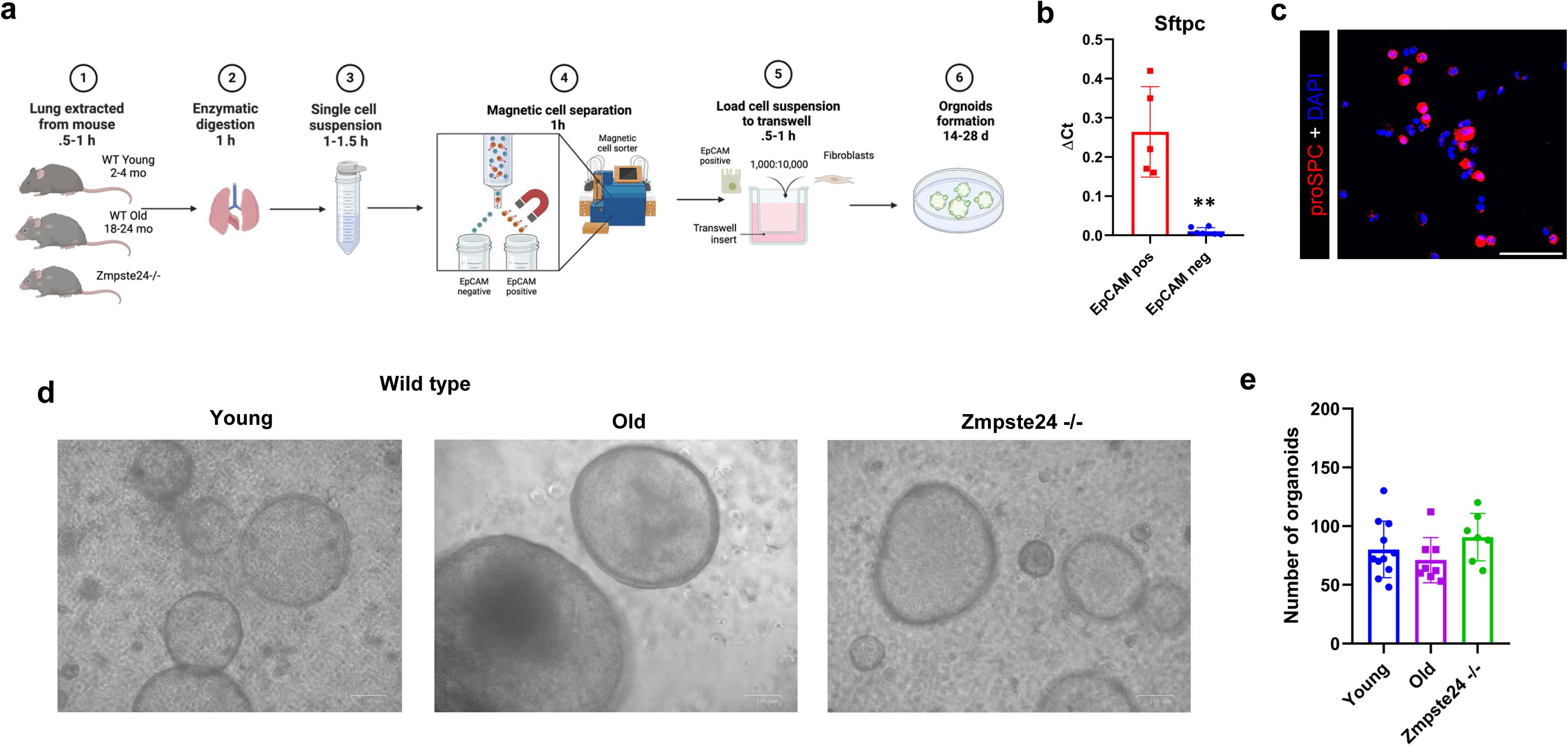
Lung progenitor cells from aged mice retain the regeneration potential. **a** Schematic representation of the experiment design. Created with BioRender.com. **b** RT-qPCR for the validation of the extraction of type 2 alveolar epithelial cells (AT2) obtained by magnetic beads (MACS) in wild-type mice by comparing the ΔCt of surfactant protein C (Sftpc) (EpCAM positive vs negative fraction). Data are presented as mean ± SEM (n = 5) by triplicated (t-test **p< 0.01). **c** Representative immunofluorescence of proSPC in EpCAM positive fraction (Scale bar, 60μm). **d** Representative brightfield micrographs of organoid cultures from wild type young, aged, and Zmpste24^-/-^ mice. (Scale bars, 100μm). **e** Graph of organoids number (n>6) Statistical analysis was performed using one-way ANOVA followed by Tukey’s post hoc test.

In this study, organoids were generated from 1,000 epithelial cells derived from young wild-type mice (12–16 weeks), physiologically aged wild-type mice (18–24 months), and Zmpste24-deficient mice, which exhibit accelerated aging (16 weeks) and with 10,000 pre-cultured fibroblasts. Our results show that isolated epithelial cells from both aging models are capable of forming organoids in numbers and sizes comparable to those generated by young wild-type mice when co-cultured with fibroblasts from young wild-type mice (**Fig. 1d and e**). An organoid growth curve was performed with old and young wild-type cells, revealing a similar trend of increase **(Supplementary Fig. 1a and b)**. In organoids derived from Zmpste24-deficient mice, proliferation was validated by Ki67 expression **(Supplementary Fig. 1c)**. After two weeks, the organoids displayed diverse multicellular structures with variations in size and morphology. As previously described, the type of organoid generated depends on the epithelial cell of origin. Larger rounded spheroids, likely corresponding to bronchiolar organoids, are characterized by smaller, more densely compacted cells, whereas alveolar organoids contain fewer cells and typically display an acinar organization **(Supplementary Fig. 1d)** [18]. Basal cells show significant pluripotency, being capable of regenerating both the airway epithelium and subsequently alveolar structures **(Supplementary Fig. 2a)** [19]. Furthermore, differentiation from AT2 into AT1 cells was confirmed by HOPX and T1α staining. These markers were observed in 14-day (SPC and HOPX) and 28-day (HOPX and T1α) organoids **(Supplementary Fig. 2b and c).**

### Autophagy is upregulated in epithelial cells from aging-accelerated mice

Since autophagy plays a critical role in stem cell maintenance, we evaluated its related gene expression in EpCAM⁺ cells isolated from freshly digested mouse lungs. A two-dimensional hierarchical clustergram was generated to show the expression profiles of 84 autophagy-related genes across young and old WT mice and Zmpste24^-/-^ groups. **Fig. 2a** shows the clustering of epithelial cells in wild-type aged mice and Zmpste24^-/-^ mice based on autophagy-related gene expression, revealing that cells from aged wild-type and Zmpste24^-/-^ mice cluster together, distinct from young wild-type cells. A volcano plot further revealed that Zmpste24^-/-^ mice cells had an increased expression of autophagy genes such as LC3, Atg12, Atg3, and Lamp1. Notably, AKt1, a negative regulator of autophagy, is found to be significantly downregulated in Zmpste24 deficient mice **(Fig. 2b).** Regarding senescence, as illustrated in **Fig 2c and d**, the EpCam+ cell fraction does not show differences in CDKN1A and p53 mRNA expression between Zmpste24^-/-^ epithelial cells and young WT cells.

**Fig. 2.**
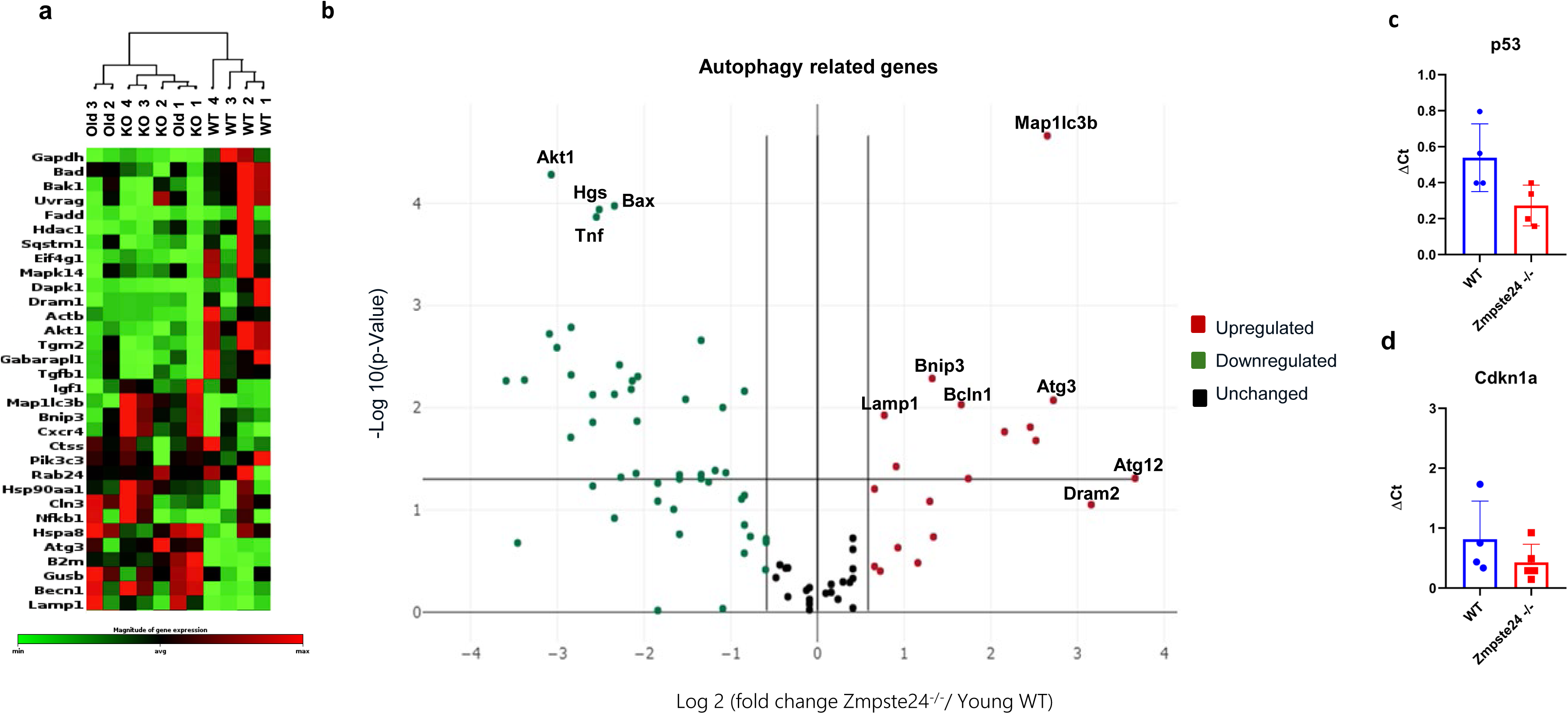
Progenitor cells show upregulation of autophagy but no changes in senescence markers. **a** Two-dimensional hierarchical clustergram of 84 autophagy-related genes analyzed using the RT² Profiler PCR Array Mouse Autophagy (PAMM-084Z, Qiagen). Samples correspond to freshly isolated EpCAM+ cells from Zmpste24^-/-^ (n=4), old wild-type mice (n=3), and young wild type mice (n=4). Fold change ≥ 2; p < 0.05. **b** Volcano plot of autophagy genes of Zmpste24^-/-^ versus young WT mice (n=4). Fold change 1.5. p-Value 0.05. **c, d** RT-qPCR for relative expression of CDKN1A and p53 in epithelial cells obtained by magnetic beads (MACS) in Zmpste24 deficient mice compared to wild-type mice (n=4).

### Senescent niche limits lung organoid growth

In the context of alveolar regeneration, strong evidence supports the notion that AT2, as facultative progenitors, require a microenvironment or "niche" that promotes such function [14,20]. Therefore, we characterized the cellular state of the fibroblasts. Aged wild-type and Zmpste24^-/-^ (>16 weeks) fibroblasts showed a significant increase in nuclear deformations, such as bleb formation, compared to young controls (**Fig. 3a and b).** Next, we evaluated senescence by SA-βgal activity, which showed a significant increase of senescent cells in Zmpste24^-/-^ fibroblasts **(Fig. 3c and d)**. Moreover, we demonstrate that in Zmpste24^-/-^fibroblasts the increase in senescence correlates with activation of the mTOR pathway, as evidenced by phosphorylation of pS6. S6 is a direct substrate of S6 kinase (S6K), which functions downstream of mTORC1, and its phosphorylation reflects activation of the mTORC1-S6K signaling axis **(Fig. 3e and f)**. By contrast, autophagy flux analyzed through LC3 and p62 after chloroquine (CQ) treatment, was decreased in these cells **(Fig. 3g and h)**, suggesting that fibroblasts reach senescence with a decrease in autophagy compared to young WT. Additionally, the senescent niche prevents organoid growth, as shown in **Supplementary Figure 3**. To confirm that Zmpste24^-/-^ fibroblasts were senescent rather than quiescent, the proliferation marker Ki67 was evaluated, revealing a lower expression of this marker in Zmpste24^-/-^ fibroblasts compared to WT cells. **(Supplementary Figure 4a and b).** In Zmpste24^-/-^ mice, fibroblast senescence is irreversible, even when treated with the pharmacological inhibitor of mTOR, Rapamycin **(Supplementary Figure 4c and d)**.

**Fig. 3.**
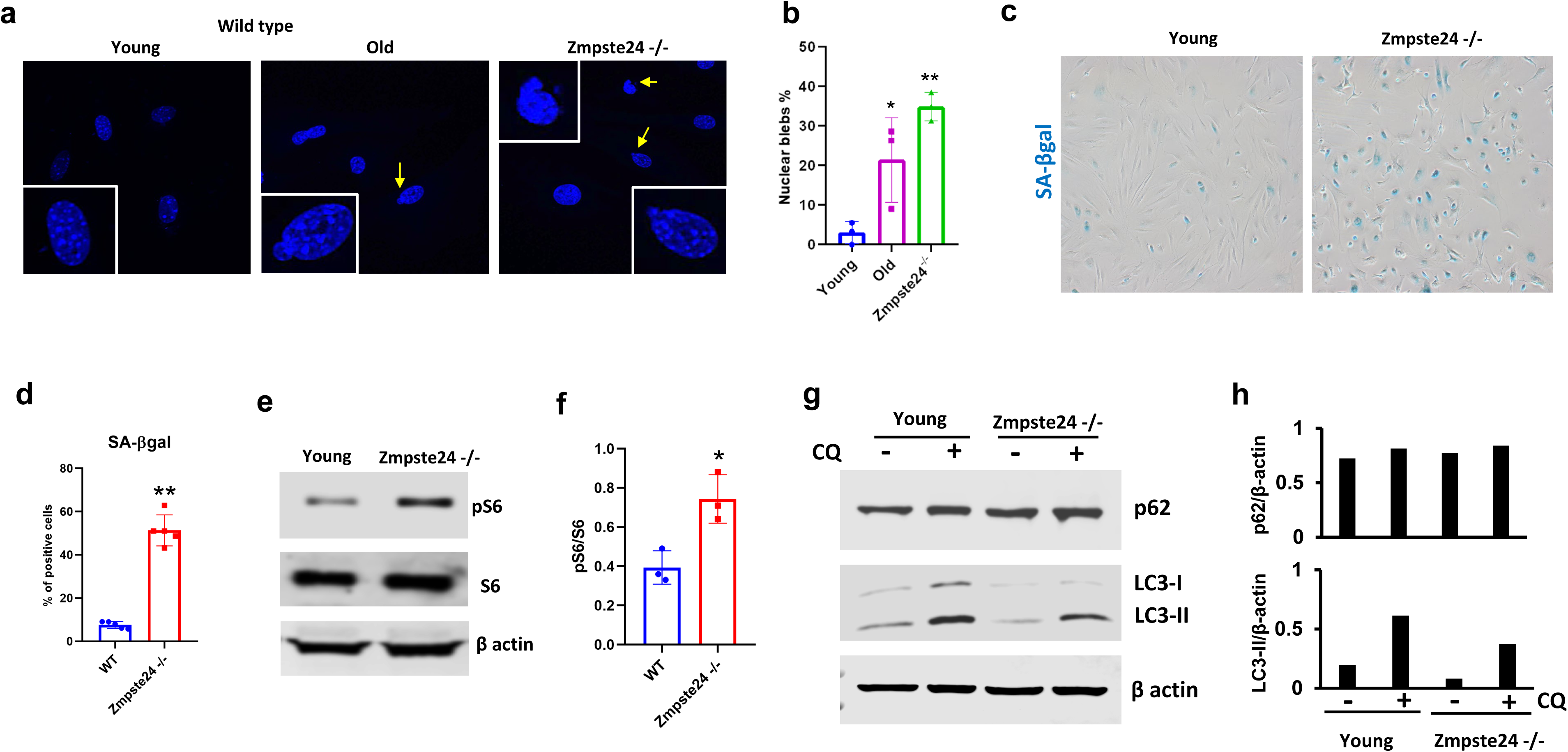
Fibroblasts from aged mice exhibit nuclear defects, early passage senescence, along with mTOR activity and reduced autophagy levels. **a** Representative image of fibroblast nuclei from young and old WT and Zmpste24^-/-^ mice. Arrowheads indicate sites of nuclear rupture. **b** Quantification of nuclei exhibiting blebs, presented as the mean percentage of nuclei analyzed from three independent animals per condition. Statistical analysis was performed using one-way ANOVA followed by Tukey’s post hoc test. **c** Fibroblasts from Zmpste24^-/-^ mice (older than 16 weeks) show significant increase of Senescence Associated -β galactosidase (SA-βgal) in fibroblasts from Zmpste24^-/-^ mice and WT at passage 3. **d** Graph of percentage SA-βgal positive cells (n=5, t-test **p< 0.01). **e** Representative immunoblot of activity of mTOR by phosphoS6 and S6 in mesenchymal cells from aged mice Zmpste24^-/-^ versus WT (β-actin as a loading control). **f** Graph of pS6/total S6 ratio (n=3, t-test *p< 0.05). **g** Representative immunoblot for autophagy marker: LC3 and p62 with/without chloroquine 10uM (CQ) for 24 hours in fibroblasts from Zmpste24 deficient mice and WT β-actin as a loading control). **h** Densitometric analysis of p62 and LC3-II levels normalized to β-actin

We induced mTOR activation in young wild-type fibroblasts through a combination of starvation and nutrient stimulation in the presence of chloroquine. This dual treatment promotes lysosomal biogenesis and changes in lysosomal acidity resulting in mTOR activation [21]. **Figure 4a** corroborates that this treatment activates mTOR signaling, as indicated by elevated levels of phosphorylated S6. Organoid culture with fibroblasts treated with this double treatment and fibroblasts from Zmpste24^-/-^ forms fewer, and smaller organoids compared to control fibroblasts (**Fig. 4b and c**). Thus, when autophagy was inhibited and mTOR activity increased, the organoids grew in a similar way as those cultured with KO fibroblasts. Furthermore, we observed senescence in these organoids. When co-cultured with wild-type fibroblasts subjected to the double-hit treatment or with Zmpste24-/- fibroblasts, the organoids also displayed senescent cells **(Fig. 4c).**

**Fig. 4.**
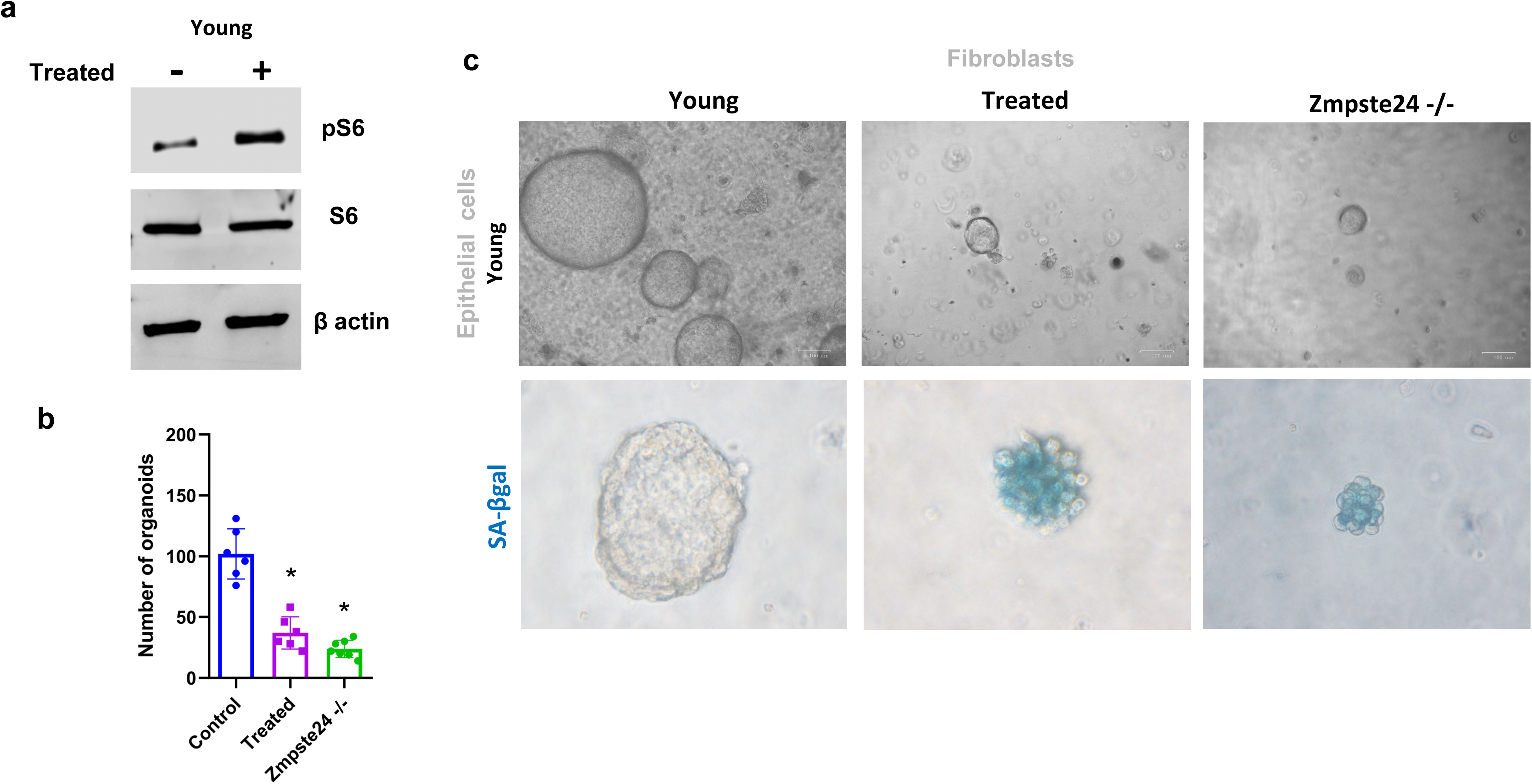
Fibroblasts from wild type mice with increased mTOR activity and from Zmpste 24 deficient mice prevent organoid formation by inducing senescence. **a** Representative Western blot show that fibroblasts from wild-type mice exhibited increased mTOR expression following sequential treatment with nutrient deprivation (HBSS, 3 h) and chloroquine (10 μM) for 24 hours **b** Number of organoids with the combination epithelial cell from wild type mice, and fibroblasts treated (HBSS 3H and CQ 10μM) for 24 hours, and from Zmpste24^-/-^ mice. Statistical comparisons were made using ANOVA test (n>6, *p<0.05). **c** Representative brightfield micrographs of organoids and senescence associated -β galactosidase (SA-βgal) stain.

### Nuclear lamins loss during lung aging

Aging is known to alter the expression of nuclear lamins, contributing to stem cell dysfunction [19,20]. However, these changes have not been characterized in the lung. In this study, we examined lamin A/C levels in lung tissues from young WT (n = 3), old WT (n = 3), and Zmpste24^-/-^ (n = 3) mice. Old mice displayed reduced lamin A/C levels throughout the lung. Quantification of fluorescence intensity, normalized to cell number, revealed a significant decrease in old WT and Zmpste24*^⁻/⁻^* mice compared with young WT mice **(Fig. 5a and b)**. Immunoblotting of nuclear fractions for lamins A/C and B1 further corroborated these findings and showed that this decline occurs gradually with age (**Fig. 5c and d**). Young mice exhibit higher expression of nuclear lamins compared to 14-month-old and 24-month-old mice, which may compromise nuclear mechanics and the tissue’s response to stress.

**Fig. 5.**
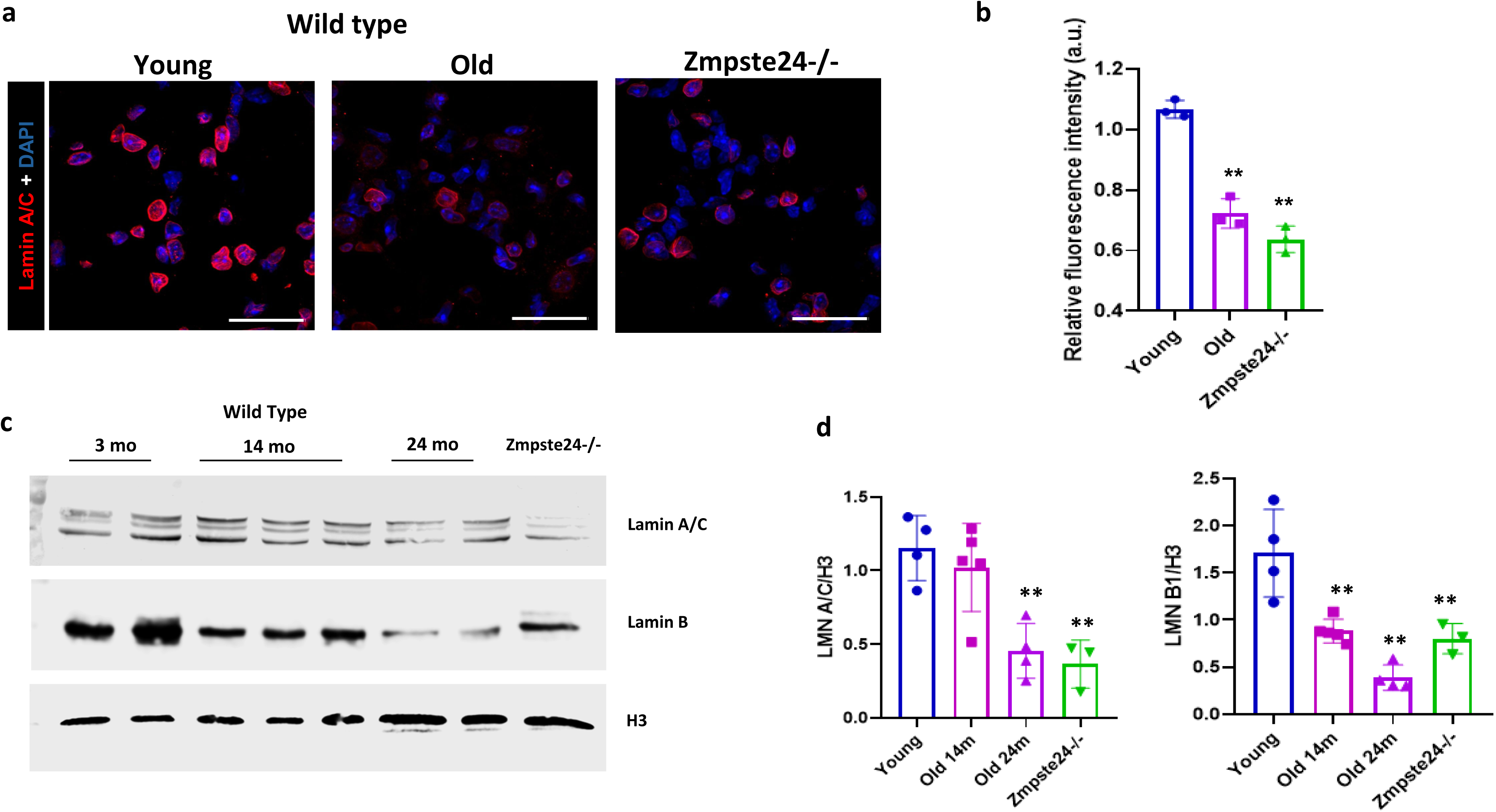
Nuclear lamins are decreased in lung tissue during aging. **a** Representative immunofluorescence images of lamin A/C in tissues from young, old, and Zmpste24^-/-^ mice (n=3 mice per group), acquired at 60X magnification with a 2.5 digital zoom. Scale bar: 20 µm. **b** Each data point in the graph represents an individual animal (n=3) and corresponds to the average immunofluorescence analysis from at least four tissue fields per animal, acquired at 40X magnification. The quantification of the mean fluorescence intensity of the nuclear lamina was normalized to the number of nuclei. Statistical analysis was performed using the ANOVA test, comparing each group to the control (**p<0.01). **c** Representative immunoblot of Lamin A/C and B1 using H3 as a loading control in young (n=3), old WT 14 months (n=3), old WT 24 months (n=2), and Zmpste24^⁻/⁻^ (n=1) mice. Band intensity was quantified and normalized to H3. **d** Graph of the densitometric analysis performed on a total sample of young (n=4), old WT 14 months (n=5), old WT 24 months (n=4), and Zmpste24^⁻/⁻^ (n=3) mice, corresponding to two independent experiments. Statistical analysis was performed using one-way ANOVA followed by Tukey’s post hoc test.

### Mechanical stress from FACS impairs epithelial cell regenerative potential

In other studies, epithelial cells from physiologically old mice have been reported to form fewer organoids [22–24]. A key difference between these studies and ours is that while previous work relied on FACS sorting to isolate epithelial cells, we used only magnetic bead-based isolation, which may influence regenerative outcomes. As illustrated in **Figure 6a and b**, incorporating an additional FACS sorting step markedly prolongs processing time and consistently reduces organoid growth under all conditions (**Fig. 6c**). Cells passing through the cytometer also express the γH2AX marker, indicating that FACS induces DNA damage (**Fig. 6e and f**). Importantly, since these cells are derived from aged mice whose mechanical properties are already compromised, this additional stress may further reduce their regenerative capacity (**Fig. 6b and d**). Although FACS is commonly used to increase cell purity, magnetic bead isolation alone yielded over 85% purity in both young and aged samples (**Supplementary Fig. 5**).

**Fig. 6.**
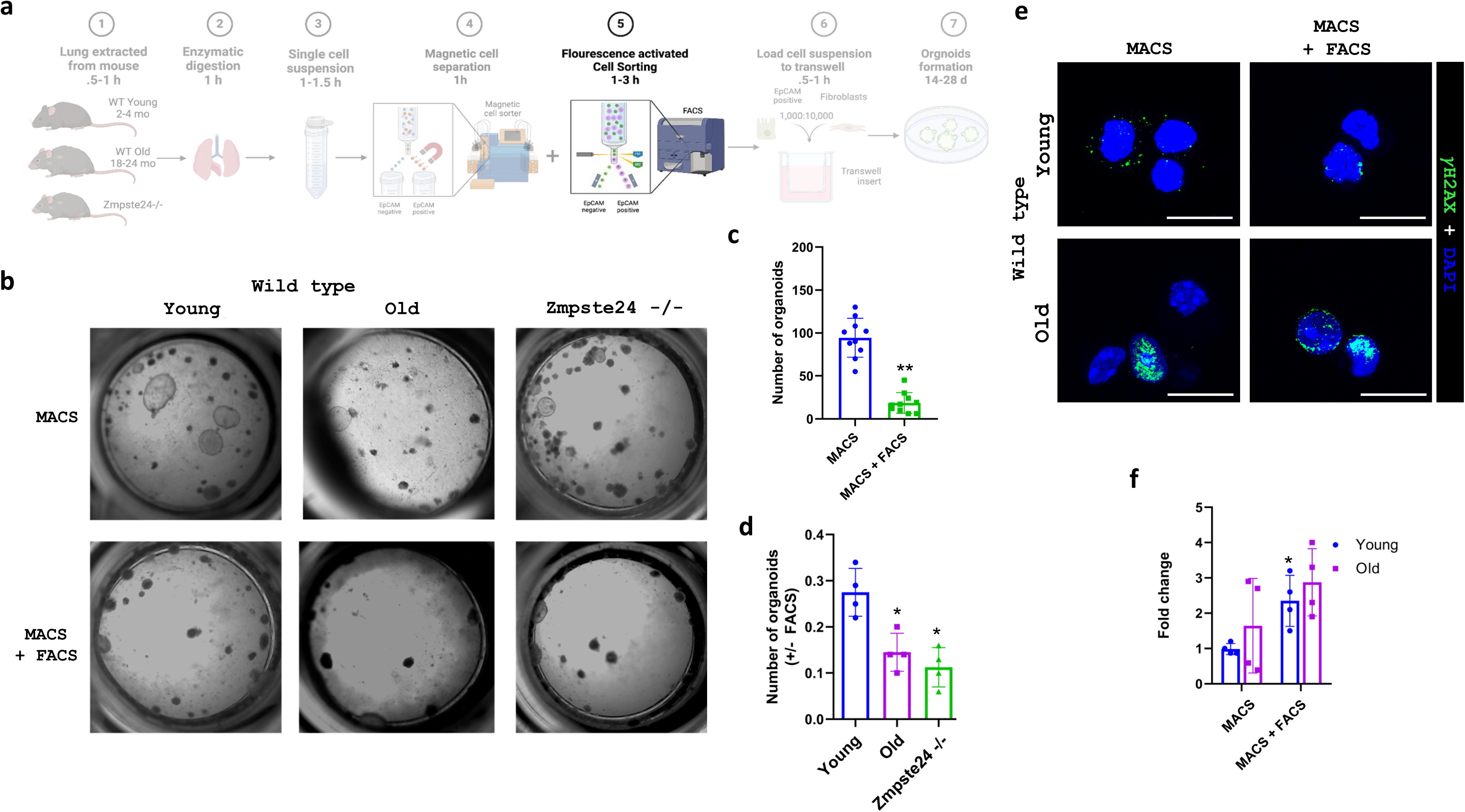
Magnetic-Activated Cell Sorting followed by Florescence-Activated Cell Sorting of epithelial cells reduces organoid growth. **a** Schematic representation of the experiment design. Created with BioRender.com. **b** Representative brightfield micrographs (Scale bars, 100μm). **c** Quantification of organoid number comparing FACS-sorted versus unsorted (control) cells (n=10, t-test **p< 0.01). **d** Graph of organoid formation by age (young (n=4), old WT 24 months (n=4), and Zmpste24^⁻/⁻^ (n=4) ratio comparing FACS-sorted versus unsorted (control) cells. Statistical comparisons were made using ANOVA test (*p<0.05). **e** Representative image of H2AX (green) immunofluorescence in EpCAM-positive cells from old mice (24 months) and young, FACS-sorted versus unsorted (control) cells. Magnification: 63X. Scale bars 10μm. **f** Quantification of γH2AX immunofluorescence signal (fold change) in EpCAM-positive cells from young and old mice, comparing FACS-sorted and unsorted (control) groups. Statistical comparisons were made using ANOVA test (*p<0.05).

## Discussion

Aging increases the morbidity and mortality associated with respiratory diseases such as chronic obstructive pulmonary disease and idiopathic pulmonary fibrosis. Contributing factors include stem cell exhaustion and altered intercellular communication, which are linked to limited regenerative capacity [9,10]. Thus, comprehending the alterations in stem cell-niche interactions is critical for understanding regeneration during aging. To investigate this, we used two models: aged wild-type mice and an accelerated-aging mouse model.

Regeneration was assessed using three-dimensional lung organoid cultures, which mimic the cell interactions of the lung and provide a platform to assess regenerative capacity [18]. Our results demonstrate that lung epithelial cells from both aged and Zmpste24^-/-^ mice maintain regenerative capacity. To investigate the mechanisms underlying this preserved function, we further characterized the cellular state of these epithelial cells. We found that epithelial cells are not senescent and maintain their lung progenitor phenotype, displaying increased autophagy, assessed through gene expression analysis. We speculate that these cells may have developed an adaptive mechanism in which autophagy contributes to maintaining a microenvironment favorable for stem cell homeostasis. Paradoxically, in Zmpste24-deficient mice, increased autophagy has been described as a systemic feature; however, this elevation does not delay aging in other organs as it appears to do in the lung [25]. Consistent with this, our cluster analysis revealed that genes related to autophagy exhibit a similar expression pattern in Zmpste24^-/-^ and aged WT mice, suggesting that this response may be a common feature of physiological aging. This convergence could be explained, at least in part, by the intrinsically low cell turnover of the lung parenchyma [26]. Moreover, *Zmpste24*-deficient mice have been reported to exhibit a protective response to bleomycin-induced fibrosis [27]. Taken together, these findings support the notion that aging may trigger a hormetic response in the lung, characterized by resistance to senescence and activation of autophagy.

Age-related decline in the lung cannot be explained solely by stem cell exhaustion but also involves pro-aging factors originating from the niche [28]. While epithelial cells in the lung encompass multiple progenitor populations, in the alveolar context our focus is on alveolar epithelial type 2 (AT2) cells and their specific niche formed by adjacent fibroblasts. These facultative progenitors play a central role in maintaining alveolar homeostasis and regeneration through close epithelial-mesenchymal interactions[6]. Our findings show that fibroblasts derived from physiologically aged and Zmpste24^⁻/⁻^ mice exhibit nuclear defects and a senescent phenotype characterized by impaired autophagy and sustained activation of mTOR signaling. When co-cultured with epithelial cells, organoid formation was markedly impaired. These findings are consistent with those reported by Chanda *et al.*, who found that fibroblasts from 24-month-old mice limit epithelial organoid growth [23]. In the same line, in human lung fibroblasts, we previously reported a persistent activation of mTOR, and a decreased autophagy associated with aging [29]. While mTOR activation has been implicated in promoting senescence, its causal role in aging within the alveolar niche remains to be established [30,31]. Additionally, pharmacological inhibition with rapamycin was insufficient to reverse fibroblast senescence, highlighting the need for additional genetic or pharmacological inhibition studies.

In this study, we also demonstrated that the expression of nuclear lamins A/C and B1 progressively decreases in the lung during both physiological aging and in the accelerated aging model Zmpste24^⁻/⁻^. This finding is relevant, as nuclear lamins are essential for maintaining nuclear structural integrity and resisting deformation; consequently, their reduction increases vulnerability to mechanical stress. In accelerated aging models such as Hutchinson–Gilford progeria syndrome (HGPS), alterations in lamin A have been shown to induce defects in chromatin organization, loss of perinuclear heterochromatin, and increased DNA damage [32]. Furthermore, reduced lamin A/C expression has been associated with nuclear reorganization characterized by the loss of lamin-associated heterochromatin domains (LADs), alterations in epigenetic marks, and disruption of three-dimensional genome architecture, with direct consequences for gene expression and tissue regeneration [33,34]. Similarly, our results suggest that the progressive loss of nuclear lamins in the aged lung may compromise regenerative capacity [32,35].

Although this link between lamin loss and progenitor cell status has not been previously studied in the lung, our findings provide initial evidence supporting a connection. We observed that the inclusion of a FACS step not only prolonged processing time but also induced DNA damage (γH2AX) and significantly reduced organoid formation, with this effect being more pronounced in aged cells, in which lamin loss already compromises nuclear recovery capacity. These results are consistent with the phenomenon described as sorter-induced cell stress (SICS), which encompasses shear forces, pressure fluctuations, abrupt accelerations, and electrical charges during sorting, all of which exceed the mechanical tolerance of cells [36–41]. When comparing cell isolation methods, we found that MACS yielded highly pure cells which displayed a better preserving viability and organoid-forming capacity, whereas FACS, although offering higher resolution, compromised cell functionality due to procedure-associated mechanical stress. Although the efficacy of MACS depends on proper optimization of antibody and bead labeling, it generally represents a faster and less damaging alternative for isolating cells [42]. Taken together, our findings demonstrate that the reduction of nuclear lamins during lung aging increases susceptibility to mechanical stress, and that technical factors such as the use of FACS can exacerbate this vulnerability. These results underscore the importance of standardizing isolation methodologies in lung organoid studies and highlight the need to further investigate lamin–chromatin dynamics and their interaction with mechanical stimuli.

A fundamental question in the biology of aging is why aging progresses at different rates within the same organism, affecting organs, tissues, and cells unequally. In the adult lung, for example, it remains unclear how the number and functional capacity of adult stem cells change with age [43]. In one of the models we used, aging is caused by a defect in the nuclear lamina, like progeria syndrome, with the phenotype developing gradually over time [8]. In our study, we have corroborated that despite having Zmpste24 deletion (at 16 weeks of age), freshly isolated Zmpste24^-/-^ cells are capable of growth like the wild type. These results confirm that aging, even linked to a genetic defect, depends on the development of specific mechanisms. Therefore, it is necessary to consider the influence of external factors such as matrix stiffness [44] particularly in aging, where mechanical stress may exacerbate nuclear alterations. In this regard, it has been proposed that during the process defined as *fibroaging*, cells undergo changes in the cellular microenvironment, receiving mechanical stimuli such as extracellular matrix stiffness, which leads to a reorganization of the cytoskeleton and induces changes in nuclear architecture, resulting in epigenetic reprogramming [45]. ECM changes have been considered as a new integrative hallmark of aging [10]. In WT mice aging is associated with the upregulation of several ECM proteins. Interestingly, although fibrillar collagen genes are not up upregulated in the Zmpste24 deficient mice, lung collagen content shows a similar increase with aging suggesting that the accumulation of this protein characteristic of aging occurs despite lamin A defects [27]. However, a potential limitation of our study is that Zmpste24^-/-^ although recapitulates key features of premature aging, it does not fully represent the complex and multifactorial nature of physiological aging.

In summary, these findings support a close interplay between progenitor cells and their niche, in which microenvironmental signals coordinate an autophagy-driven stress response to promote regeneration. Understanding the mechanisms by which these cells lose function with aging will provide deeper insight into their role in the pathophysiology of various respiratory diseases, which could lead to the development of more effective therapeutic strategies.

## Methods

### Murine model

Animal use complied with relevant guidelines and regulations for animal care, and the protocol was approved by the Bioethics Committee of the Instituto Nacional de Enfermedades Respiratorias “Ismael Cosio Villegas” (Approval Codes: B01-2023 and B17-25). C57BL/6J mice (wild type) of 12-16 weeks and 18–24 months were used as young and physiologically aged groups, respectively. Zmpste24^-/-^ mice were used as a model of accelerated aging; animals were included from 16 weeks of age, and both genotyping and phenotype characteristics were confirmed prior inclusion **(Supplementary Fig. 6).** This strain was developed and kindly donated by Dr. Carlos López-Otín.

### Isolation of lung epithelial cells and fibroblasts

Lungs of WT and Zmpste24^-/-^ mice were perfused with PBS 1X and instilled with dispase (Corning # 47743) for 45 min at room temperature. Trachea and upper airways were removed. The lung was mechanically dissociated and filtered through 70μm and 40μm nylon filters. After Red Blood Cell Lysis Buffer was used. The cell suspension was incubated for epithelial cell positive selection with anti-mouse CD326 (EpCAM) attached with magnetic beads (Miltenyi Biotec #: 130-105-958). Epithelial cells were used immediately, and purity was determined by RT-qPCR of surfactant protein C (SPC). Lung fibroblasts cells were obtained from the negative fraction. Cells were plated on, and media was changed 24h after to remove cells that were not attached (non-mesenchymal). Fibroblasts were maintained in culture with DMEM medium with 5% FBS in a 37°C incubator in an atmosphere of 95% air and 5% CO2 until use. Cells were starved in HBSS (Gibco, #14025092) for 3 hours, then stimulated with complete medium for 24 hours in the presence of 10 μM chloroquine (Sigma, #C6628) to induce mTOR activation and block lysosomal degradation.

### Organoids lung culture

For lung organoid culture, freshly harvested epithelial cells (EpCam positive) were cocultured with fibroblasts; in a ratio of 1,000:10,000 (epithelial: fibroblast). Fibroblasts and epithelial cells were isolated from different mice grown in the same cages. The fibroblasts were previously cultured from different mice on preceding days and expanded up to passage 3 before being used to ensure optimal cell viability. Briefly, epithelial cells and fibroblasts were co-cultured in a 1:1 Matrigel–culture medium mixture (Corning, Cat. #356237, Lot #3061001). MEMα-based medium (Gibco, Catalog # M4526) was supplemented with 10% FBS, 1× insulin/transferrin/selenium, 2 mM L-glutamine, penicillin-streptomycin, 0.25 μg/mL amphotericin B, 0.0002% heparin and ROCK pathway inhibitor at 1µM (Selleckchem, Catalog # S1049). The 6.5 mm transwell with 0.4μm pore polyester membrane inserts (Corning, Catalog # CLS3470) were required for culture to establish an air-liquid interface. Organoids were cultured with media changes every 2 days and fixed using Fixation Buffer (Sigma, Catalog # F1797). Since organoids are distributed at varying depths in Matrigel, they were counted from a single focal plane within the z-stack, chosen based on the highest density of visible organoids. Photographs of bright fields were taken using an inverted microscope (Nikon) and images were analyzed with NIS-Elements Software.

### Senescence Associated-β galactosidase assay (SA-βgal)

Senescence associated β-galactosidase (SA-βgal) staining was performed in fibroblasts and transwell-inserts containing organoid cultures with the Senescence Cells Histochemical Staining Kit (Sigma, Catalog # CS0030) according to the manufacturer’s instructions.

### RT-qPCR

RNA of epithelial cells was isolated using Trizol reagent (Invitrogen) and cDNA was synthesized from 500ng of RNA using iScript cDNA Synthesis Kit (Bio-Rad) according to manufacturer’s protocol. Quantitative PCR amplification was performed in a mixture containing 6ng of cDNA, and the following primers: CDKN1A Rv:CCTGTTCTAGGCTGTGACTGCTT; Fwd: CATTCCCTGCCTGGTTCCTT, and p53 Rv: GGCGAA AAGTCTGCCTGTCTT, Fwd: CGCTGCTCCGATGGTGAT, β-actin Rv: GGCTGTATTCCCCTCCATCG, β-actin Fwd: CCAGTTGGTAACAATGCCATGT. In addition, TaqMan probes were used for SPC and 18s quantification. Thermocycler QS6Flex (Applied Biosystems) was used for qPCR experiments, ΔCt was used for data analysis.

Gene expression analysis of mouse autophagy-related genes was performed using the RT² Profiler PCR Array Mouse Autophagy (PAMM-084Z, Qiagen) and processed with the RT² Profiler PCR Data Analysis tool (GeneGlobe, Qiagen). cDNA samples were obtained from epithelial cells of four KO mice and four WT mice. Ct values were normalized to the arithmetic mean of the housekeeping genes (Actb, B2m, Gapdh, and Gusb).

### Immunocitochemistry

Paraffin-embedded organoid sections (3 µm) were prepared for IHC. Antigen retrieval was performed using a citrate buffer, followed by permeabilization and blocking. Sections were incubated overnight at 4 °C with the primary antibody against proSPC (Millipore, Catalog #AB3786). After washing, samples were incubated with the appropriate secondary antibody, followed by horseradish peroxidase–conjugated streptavidin, and counterstained with hematoxylin. Images were acquired using an inverted microscope (Eclipse, Nikon) and analyzed with NIS-Elements software.

### Immunofluorescence

Lungs were perfused with 1X PBS to remove blood, fixed with 10% neutral buffered formalin (NBF), and embedded in paraffin blocks. Antigen retrieval was performed using a citrate buffer, followed by permeabilization and blocking. Sections were incubated overnight at 4 °C with the primary antibody against Lamin A/C (GeneTex, Catalog #GTX101127). After primary antibody incubation, Alexa Fluor 546 donkey anti-rabbit secondary antibody (Invitrogen, Catalog #A10040) was applied. Nuclei were counterstained with DAPI (Sigma Catalog #D1306). EpCAM-positive cells were seeded on coverslips coated with Matrigel and incubated for 2 h. Cells were then fixed with 4% paraformaldehyde for 20 min at room temperature, followed by two washes with 1X PBS. Blocking and permeabilization were performed with 0.5% Triton X-100 in PBS containing 5% BSA for 45 min. Cells were incubated overnight at 4°C with the following primary antibodies: SPC (Millipore, Catalog #AB3786), HOPX (Santa Cruz Biotechnology, Catalog # sc398703), Anti-Podoplanin/ T1alpha Syrian hamster monoclonal antibody (Abcam, # ab11936), Ki67 (Abcam, Catalog # ab833), and H2AX (Abcam, Catalog #ab11174). The corresponding secondary antibodies were Alexa Fluor 488 donkey anti-rabbit (Invitrogen, Catalog #A-21206) or Alexa Fluor 549 donkey anti-mouse (Invitrogen, Catalog #A-21203). Nuclei were counterstained with DAPI (Catalog #D306). Images were obtained using confocal Leica TCS-SP8.

### Western Blot

Proteins were extracted from cells using RIPA buffer lysis (Sigma Catalog #R0278), and 20μg of total protein was used in polyacrylamide gels for electrophoresis. Nitrocellulose membrane was used to transfer proteins and block for 1 h. Primary antibodies for LC3 (Sigma, Catalog #L7543), P62 (Abcam, Catalog #ab56416), S6 (CST, Catalog #2217), p(235/236)S6 (CST, Catalog #2211) Lamin b1 (Invitrogen, Catalog #702972), Lamin A/C (GTX, Catalog #101127) and β-actin (Sigma, Catalog #A5441) or H3 (GTX, Catalog #122150) as a loading control were incubated overnight. Finally, Li-Cor infrared secondary antibodies and Odyssey scanner (Li-Cor) were used. Quantification was done using ImageJ software (NIH).

### Flow cytometry

After separation with magnetic beads, Ep-CAM-positive cell suspensions were stained with anti-CD326 PE-Cy7 antibody (BioLegend) for 30 minutes at 4°C, avoiding light. They were subsequently washed and subjected to fluorescence-activated cell sorting. Using a FACS Aria II (BD), cells were identified by gating singlets, and CD326+. Double-selected (isolated with magnetic beads and FACS) Ep-CAM-positive cells were cultured to obtain organoids.

### Statistical analysis

Data are presented as mean ± standard deviation. When the assumptions of normality and homogeneity of variances were satisfied, statistical comparisons were performed using either a two-tailed Student’s *t*-test (for comparisons between two groups) or one-way analysis of variance (ANOVA) followed by Tukey’s post hoc test (for comparisons among more than two groups). A *p*-value of less than 0.05 was considered statistically significant. All statistical analyses were conducted using GraphPad Prism, version 9.

## Author Contributions

Y.R. conceived and designed research; Y.R., F.T-M., A.G.-V., U.C-S., T.V-H., R.R., E.H-X. and M.R.L. performed experiments and analyzed data. Y.R. and F.T-M. drafted the manuscript; M.S. and A.P. evaluated the results and edited and revised the manuscript. All authors approved the final version of the manuscript.

## Acknowledgments

We would like to express our gratitude to Edgar Jiménez for all the technical suggestions. We also thank, Guadalupe Hiriart Valencia, Erika Liliana Monterrubio Flores, Alberto Pizaña Venegas, and the biotherium staff at INER for their technical support. Additionally, we appreciate the support from the Laboratorio Nacional Conahcyt de Investigación y Diagnóstico por Inmunocitofluorometría (LANCIDI) and Laboratorio Nacional de Soluciones Biomimeticas para Diagnóstico y Terapia (LaNSBioDyT).

## Conflict of interest

None declared.

## Ethical Statement

All experimental procedures were conducted in accordance with the ethical standards of the Comité de Ética en Investigación, Comité de Bioseguridad, and Comité Interno para el Cuidado y Uso de Animales de Laboratorio (CICUAL) of the Instituto Nacional de Enfermedades Respiratorias Ismael Cosío Villegas, and were approved under protocol numbers B01-23 and B17-25.

## Funding

The present study was supported by SECIHTI Ciencia de Frontera 2019 #51219.

## Data availability statement

All data that support the findings of this study are available from the corresponding author upon reasonable request.

## Legends of supplementary figures

**Supplementary Fig. 1 Lung organoids growth from aged mice**

**a** Quantification of organoid growth by age (young, *n* = 4; old WT, 24 months, *n* = 4). Statistical significance was determined using one-way ANOVA.

**b** Brightfield micrographs showing organoids derived from 24-month-old WT mice at different days (d) of culture. Scale bar: 100 μm.

**c** Immunofluorescence staining for Ki67 in 14-day-old organoids derived from Zmpste24^⁻/⁻^ mice at 16 weeks. Scale bar: 20 μm

**d** Representative brightfield micrograph of alveolar (top) and airway (bottom) in 14-day-old organoids derived from Zmpste24^⁻/⁻^ mice. Scale bar: 50 μm.

**Supplementary Fig. 2 Lung organoids characterization from aged mice**

**a** Immunofluorescence staining for SPC in 14-day-old large basal organoids. Scale bar, 30 μm.

**b** Staining for SPC and HOPX in 14-day-old organoids illustrating initial alveolar differentiation. Scale bar, 20 μm.

**c** Staining for T1α and HOPX in 28-day-old organoids showing alveolar type I differentiation. Scale bar, 50 μm.

**Supplementary Fig. 3 Organoids formation by epithelial cells and fibroblast from Zmpste24^⁻/⁻^ mice**

**a** Representative brightfield micrographs (Scale bars, 100μm)

**b** Graph of organoids number.

**Supplementary Fig. 4 Reduced proliferation and persistent senescence in Zmpste24^-/-^fibroblasts upon rapamycin treatment**

**a** Representative immunofluorescence images of Ki67 (red) in young WT and Zmpste24^-/-^fibroblasts counterstained with DAPI (blue).

**b** Bar graph showing quantification of mean nuclear fluorescence intensity in young WT and Zmpste24^-/-^ fibroblasts from three independent experiments (t-test, p < 0.01).

**c** Fibroblasts from Zmpste24*^⁻/⁻^* mice (older than 16 weeks) show a significant increase in SA-β-gal activity compared with WT fibroblasts at passage 3, which was not reversed by 24 h treatment with 20 nM rapamycin.

**d** Graph of percentage SA-βgal positive cells (n=3).

**Supplementary Fig. 5 Purity Analysis by Cytometry in EpCAM positive cell suspensions** Flow cytometry gating strategy. The dot plots show the subsequent gates for analysis of (EpCam)-positive population. Forward scatter area (FSC-A) and forward scatter height (FSC-H) parameters were applied to exclude cell doublets. Subsequent plots were generated to identify EpCam-positive cells. The purity of each cell population is reported as a percentage of the total cell count. **a** Young gating strategy. **b** Old gating strategy.

**Supplementary Fig. 6 Genotyping of Zmpste24 deficient mice**

**a** Photograph of young wild-type and Zmpste24 mice at 16 weeks.

**b** Representative gel electrophoresis of PCR with specific primers for Zmpste24 WT and KO.

## Notes

### Competing Interest Statement

The authors have declared no competing interest.

