## Supplementtary figures for "Lung progenitors’ exhaustion in response to microenvironment stress during aging"

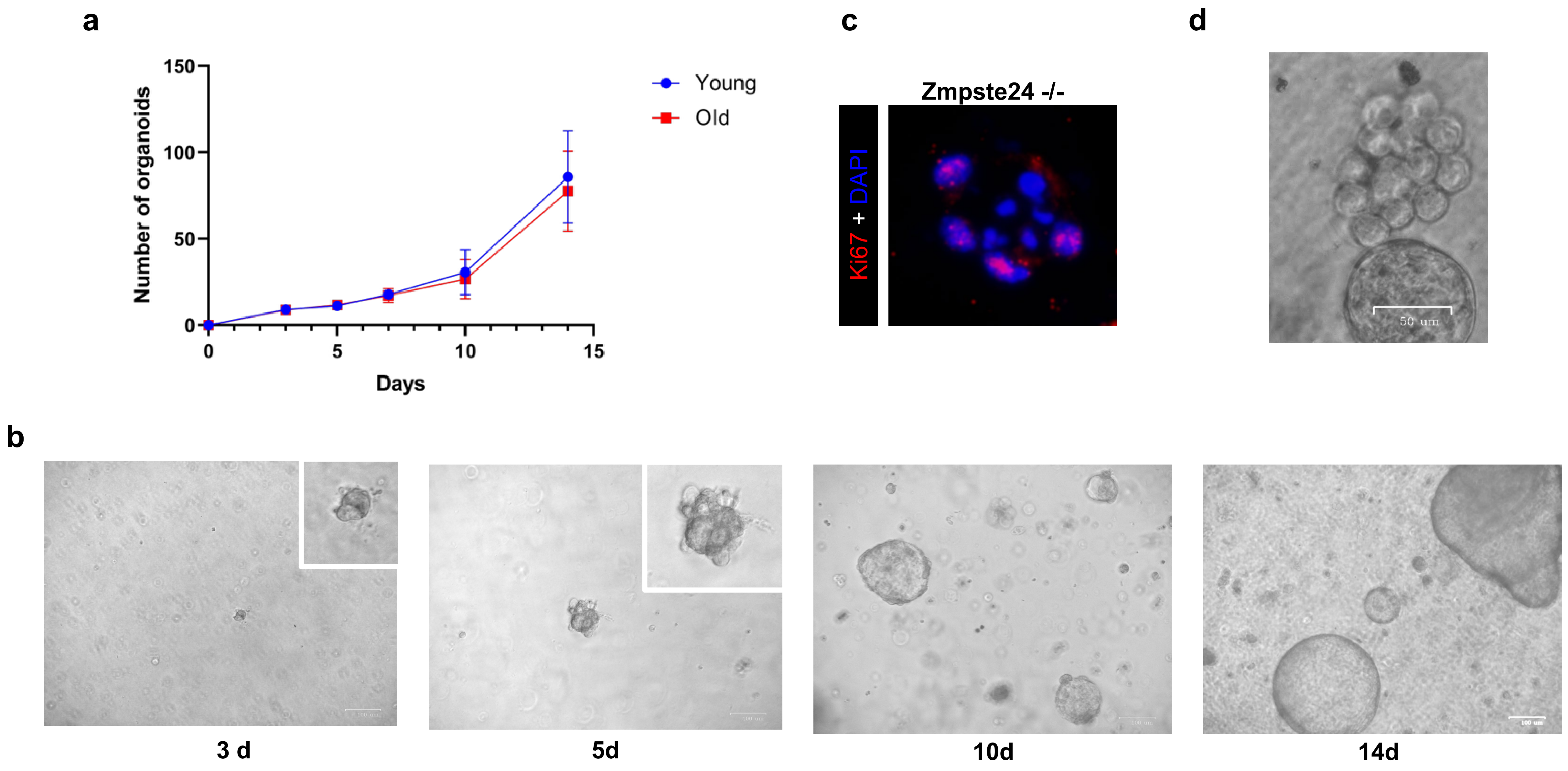


**Supplementary figure. 1**

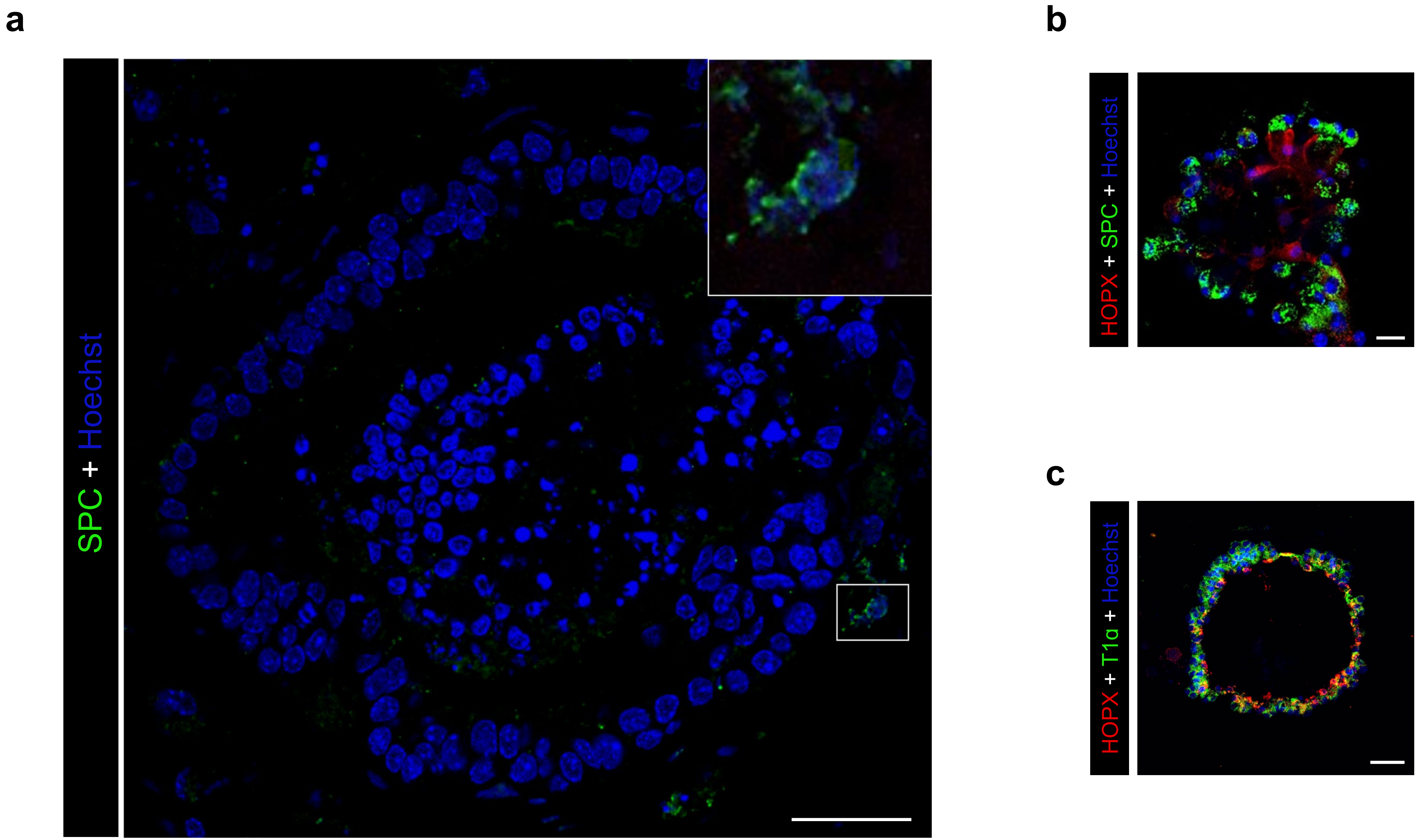


**Supplementary figure. 2**

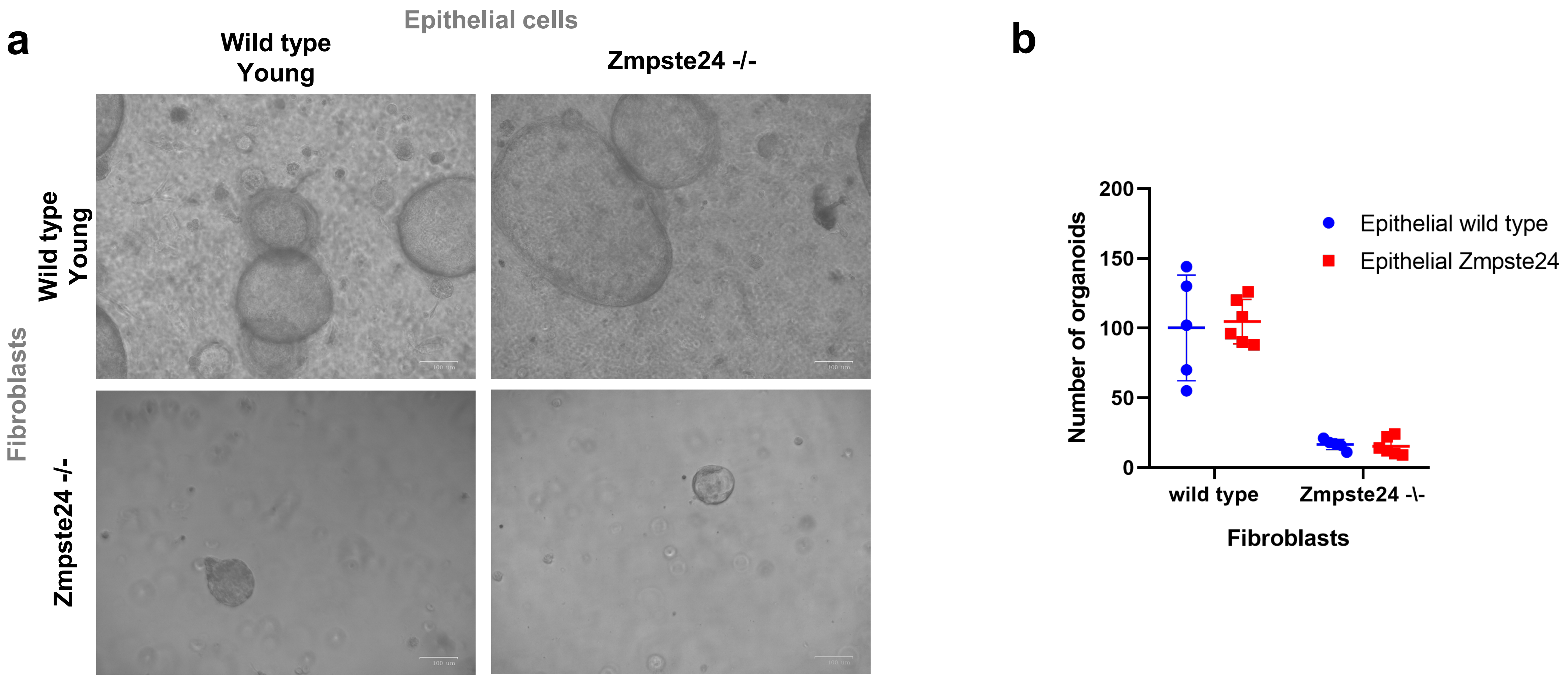


**Supplementary figure. 3**

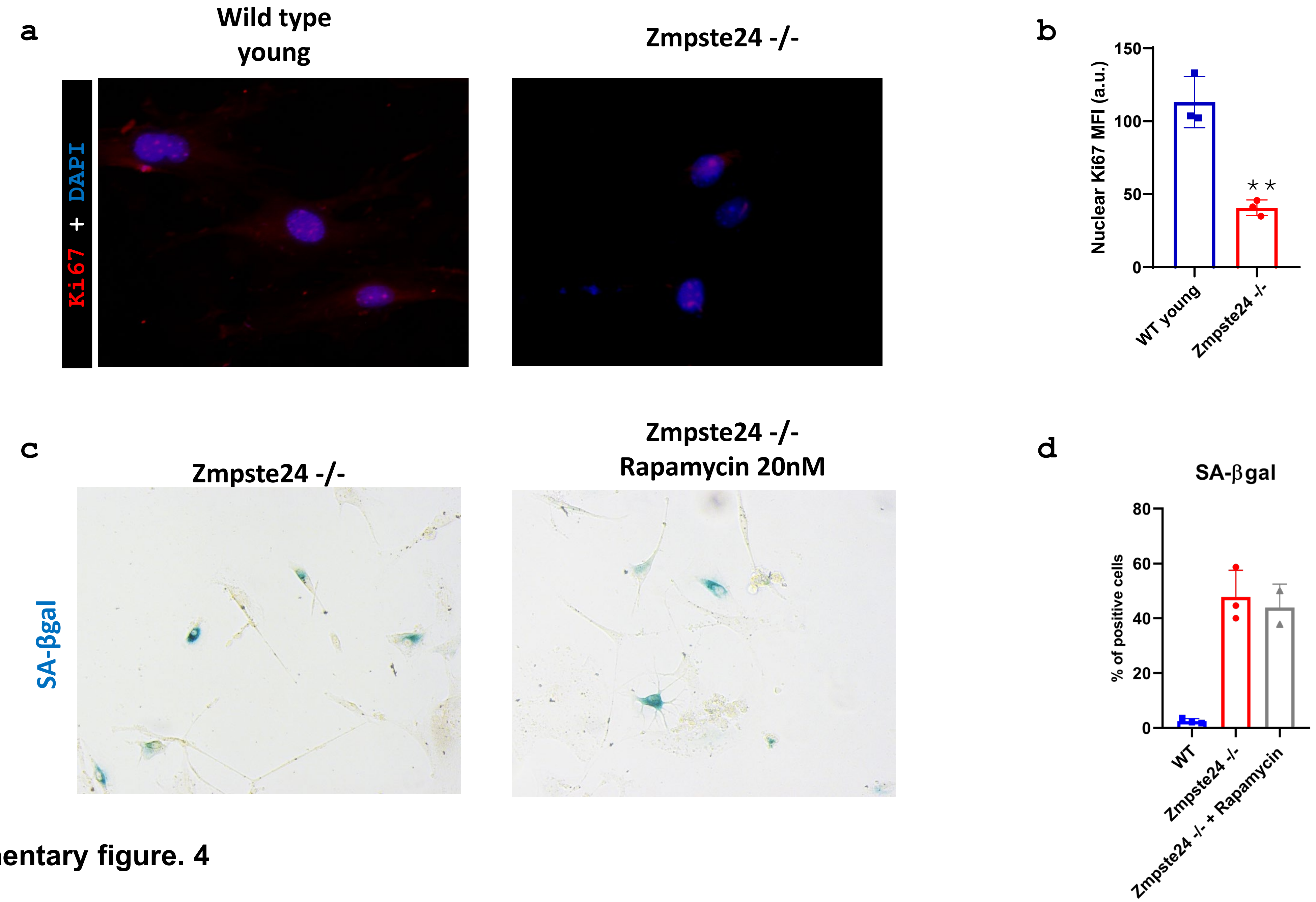


Supplementary figure. 4

**a**

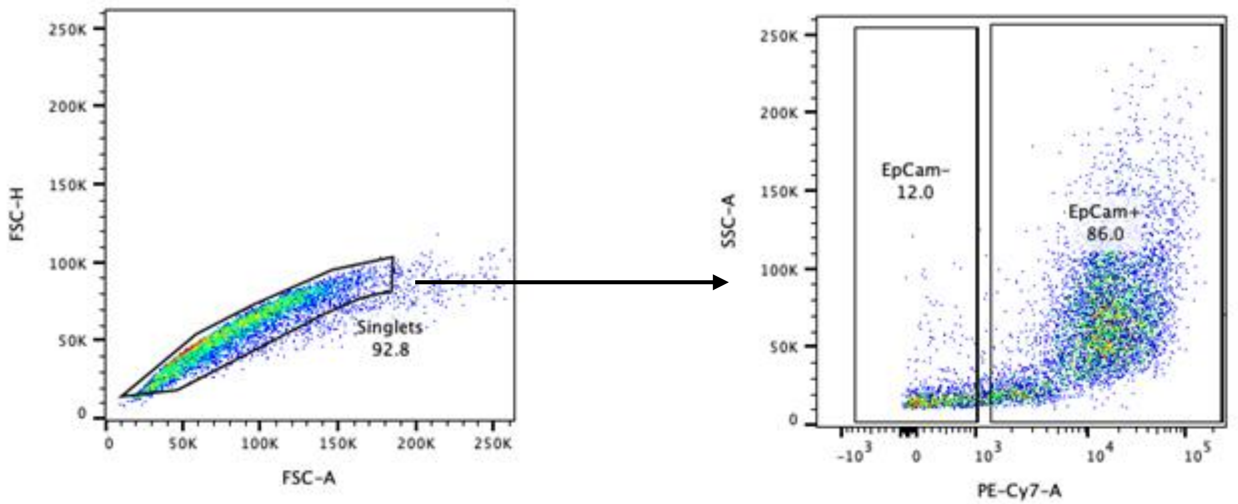


**Young**

**b**

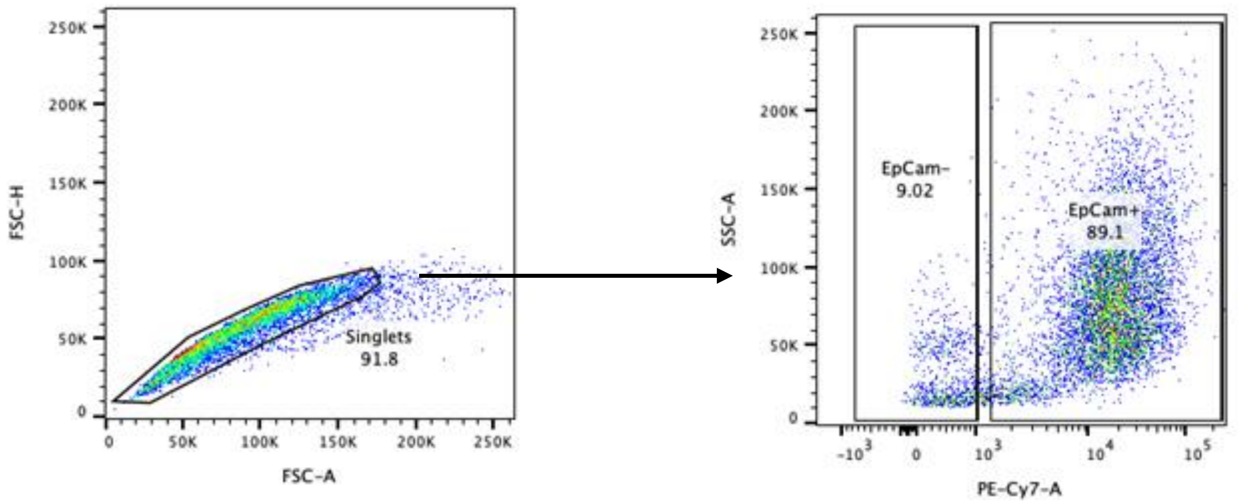


**Old**

**Supplementary figure. 5**

**a**

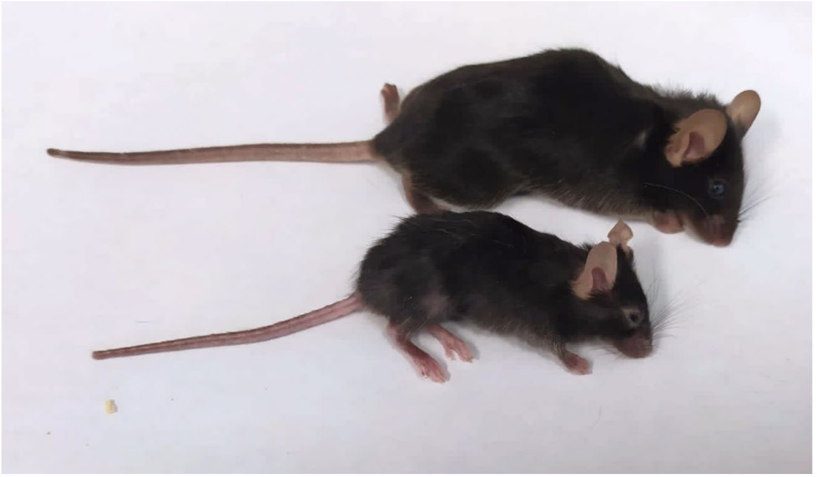

**b**

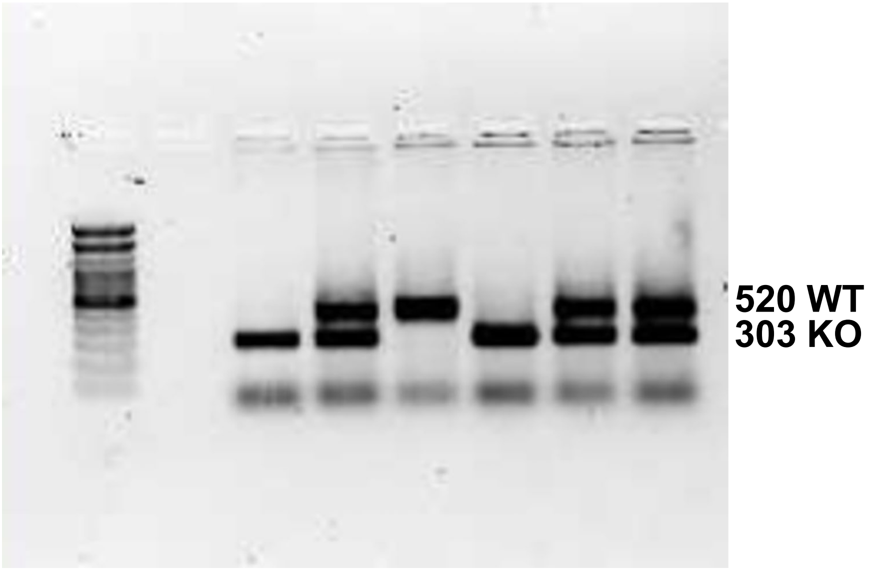


**Supplementary figure. 6**
